# A Mediator-dependent hypertranscriptional program governs neural stem cell fate decisions *in vivo*

**DOI:** 10.64898/2026.02.26.708183

**Authors:** Tiago Baptista, Daniela Lopes, Ana Rita Rebelo, Catarina CF Homem

## Abstract

Hypertranscription, a global increase in transcriptional output, is a defining feature of stem cells, but its biological relevance and regulation *in vivo* remain unclear. Using *Drosophila* neural stem cells (neuroblasts), we investigate the function and regulation of hypertranscription. Neuroblasts exhibit higher transcriptional activity than their differentiated progeny, establishing them as a model of hypertranscriptional stem cells. We identify the Mediator complex as a key regulator of this elevated transcriptional state. Mediator is broadly enriched across neuroblast chromatin, and its depletion causes a substantial, selective reduction in transcription in neuroblasts, with only mild effects in differentiated cells. Functionally, Mediator depletion impairs lineage progression, causing an accumulation of undifferentiated, stem-like cells, highlighting the necessity of hypertranscription for proper stem cell fate and differentiation. Finally, we show that *Drosophila* brain tumor neuroblasts exhibit an exaggerated, Mediator-dependent hypertranscriptional program. These findings establish Mediator as a central regulator of hypertranscription in both normal and tumorigenic neural stem cells, crucial for neural differentiation and stem cell fate progression.

## Introduction

The transition from stem cell (SC) self-renewal to differentiation is a complex process regulated by a delicate interplay of transcriptional, metabolic, and epigenetic mechanisms(Harvey et al. 2019). A defining but still enigmatic feature of SCs across diverse systems is hypertranscription, a global elevation in transcriptional output in SCs relative to more differentiated progeny in their developmental lineages. This includes housekeeping genes, lineage-specific genes, and non-coding genomic regions. Hypertranscription has also been associated with disease, particularly in cancer, where many tumors exhibit markedly increased transcriptional output compared to their non-malignant counterparts(Zatzman et al. 2022). Despite its prevalence, hypertranscription remained largely unrecognized until the advent of cell-number-normalized transcriptomics, as earlier averaged bulk-transcriptome approaches masked transcript abundance per-cell (Kim et al. 2023).

Some of the strongest evidence for hypertranscription emerged from *in vitro* mammalian studies. Naïve embryonic SCs (ESCs), which are more pluripotent, display higher hypertranscription than primed ESCs (Kim et al. 2023; Bulut-Karslioglu et al. 2018a). Mouse ESCs also display higher transcriptional output than more differentiated neural progenitors(Percharde et al. 2017), strengthening the idea that elevated transcriptional activity is tightly linked to SC potency. More recent work has extended these observations to *in vivo* contexts, including in mouse primordial germ cells(Percharde et al. 2017), hematopoietic and epidermal progenitors (Kim et al. 2023), and brain(Lavado et al. 2018a). Collectively, these findings suggest that hypertranscription may be a conserved, biologically meaningful state in stem cells. Several molecular players such the MYC family of transcription factors (TFs)(Scognamiglio et al. 2016; Nie et al. 2012), the ATP-dependent chromatin remodeler Chd1(Guzman-Ayala et al. 2015; Gaspar-Maia et al. 2009; Bulut-Karslioglu et al. 2021), and the YAP/TAZ transcriptional regulators(Lavado et al. 2018a) have been implicated in assisting hypertranscription amplification or maintenance both *in vitro* and *in vivo*. Yet, despite these insights, two major questions remain unanswered: what molecular mechanisms initiate or extinguish global transcriptional amplification in SCs, and what is the *in vivo* functional relevance of sustaining such elevated transcriptional states in SCs?

Clues may lie in the broader architecture of SC transcriptional networks. The core gene expression program governing mammalian ESCs identity relies on a small set of master TFs(Ng and Surani 2011; Orkin and Hochedlinger 2011; Young 2011) that act together with the Mediator complex, a conserved transcriptional coactivator that bridges enhancers to RNA Polymerase II (Pol II) to activate and maintain stem cell regulatory circuits (Borggrefe and Yue 2011; Conaway and Conaway 2011; Kornberg 2005; Malik and Roeder 2010a; Kagey et al. 2010). Notably, Mediator occupancy has been used to rank enhancers in ESCs, with those exhibiting the highest Mediator occupancy coined as super-enhancers(Whyte et al. 2013), a class of enhancers particularly enriched for TFs motifs via which Mediator can regulate the expression of key genes in cell identity and oncogenes(Hnisz et al. 2015). During differentiation, Mediator is redistributed to lineage-specific enhancers to rewire gene expression programs and drive cell fate transitions(Whyte et al. 2013). These properties position Mediator as a compelling, though untested, candidate regulator of hypertranscription. Supporting this notion, CDK8/19 inhibition, which stimulates Mediator activity, is sufficient to repress differentiation and promote self-renewal of mouse ESCs in a more naïve state, characterized by higher levels of transcription compared to the primed state(Lynch et al. 2020). Mediator is also essential for development in diverse organisms, as loss of its components results in severe defects or embryonic lethality(Miao et al. 2018; Rocha et al. 2010; Westerling et al. 2007; Ito et al. 2000). In neurogenesis, Mediator has been shown to be necessary for terminal differentiation of *Drosophila* neural SCs (NSCs)(Homem et al. 2014), while loss of Mediator impairs the formation of newborn neurons in the subgranular zone of the adult mouse hippocampus(Chen et al. 2020). Similarly, impairment of Mediator activity in zebrafish resulted in defective brain development and cerebellar atrophy(Li et al. 2025), phenocopying the human neurodevelopmental syndrome caused by biallelic variants in *MED27*(Meng et al. 2021). Given Mediator’s central position in transcriptional regulation and its established role in SC maintenance and fate transitions, it is a strong candidate for orchestrating hypertranscription within developing lineages, yet this possibility has not previously been explored.

To investigate the *in vivo* role of hypertranscription in fate decisions and differentiation, and to test the hypothesis that Mediator regulates this process, we turned to *Drosophila melanogaster* NSCs – the neuroblasts (NBs). NBs generate all neurons and glia that populate the animal brain, and both NBs and their differentiated progeny can be precisely tracked and genetically manipulated *in vivo*(Homem and Knoblich 2012), making this system ideally suited for mechanistic dissection of hypertranscription. By the combinatorial use of transcriptomics, recruitment studies, imaging, and biochemical assays, we show that NBs have higher transcriptional output than their more differentiated progeny, demonstrating that hypertranscription is conserved in *Drosophila.* Importantly, the elevated transcriptional output of NBs cannot be explained simply by differences in cell size. We further show that the Mediator complex is a key regulator of this hypertranscriptional state: it is broadly spread throughout chromatin in NBs, and its depletion results in abrupt NB-specific decrease in transcription, with only mild effects in differentiating or differentiated progeny. This NSC selective sensitivity establishes Mediator as a selective and *bona fide* regulator of hypertranscription, unlike general transcriptional regulators which would be expected to impair transcription uniformly across lineages. Additionally, abrogation of Mediator-dependent hypertranscription from NBs disrupts neural lineage progression, resulting in the accumulation of NB-like cells at the cost of more differentiated cells, ultimately impairing neurogenesis. Given Mediator’s known involvement in metabolic regulation in NSCs(Homem et al. 2014), we also examined whether metabolism contributes to hypertranscription. While Mediator loss alters NB metabolic state, metabolic perturbations alone do not impair hypertranscription, indicating that hypertranscription is not simply a downstream consequence of metabolic activity. Finally, we also show that NB-derived brain tumors display an exaggerated hypertranscriptional program that is likewise Mediator-dependent, underscoring the relevance of transcriptional amplification to both normal development and disease. Altogether, these findings identify Mediator as a master regulator of hypertranscription in NSCs and uncover its essential role in linking transcriptional amplification to fate transitions and tumor progression.

## Results

### Neuroblasts exhibit hypertranscription as determined by elevated transcriptional output

Although hypertranscription has been described in several models, it has never been reported in *Drosophila melanogaster*. To investigate the existence of hypertranscription in *Drosophila*’s NSCs relative to their differentiated progeny, we analyzed previously published single-cell RNA sequencing (scRNAseq) data from the central brain (CB) of wandering third instar larvae, including both type I (NBI) and type II (NBII) NBs and their lineages. NBI divides asymmetrically to self-renew and generate a more fate-committed ganglion mother cell (GMC), that divides symmetrically to originate NBI-derived neurons or glial cells. In contrast, NBII divide asymmetrically to produce an intermediate neural precursor (INP) instead of a GMC. Following a maturation phase, INPs undergo several rounds of asymmetric division, self-renewing and generating a GMC, which then divides to originate two differentiated cells(Homem and Knoblich 2012). The analyzed scRNAseq dataset captures all these different cell types and includes comprehensive annotations based on the expression patterns and average levels of established marker genes. 39 distinct clusters were identified in this dataset, that can be combined in larger groups corresponding to NBIs, NBIIs, INPs, GMCs and early-born neurons(Marques et al. 2023). Since hypertranscription can be characterized both qualitatively (by the number and type of genes expressed) and quantitatively (by the total number of transcripts per cell), we assessed both the number of detected features (genes) and total unique molecular identifiers (UMIs) counts across the cell clusters. NBIs and NBIIs displayed the highest number of expressed genes (average of 4016 and 3834 features, respectively), followed by INPs, type I and type II GMCs and, lastly, neurons (average of 1996, 1569, 1149, and 1283 features, respectively) (Figure 1A). Similarly, NBIs and NBIIs presented the highest average counts per cell (39427 and 37205 counts per cell), followed by INPs, type I and type II GMCs, and neurons (average of 8733, 5696, 3321, and 3528 features, respectively), suggesting that NBs are transcriptionally more active than their progeny (Figure 1A).

**Figure 1.**
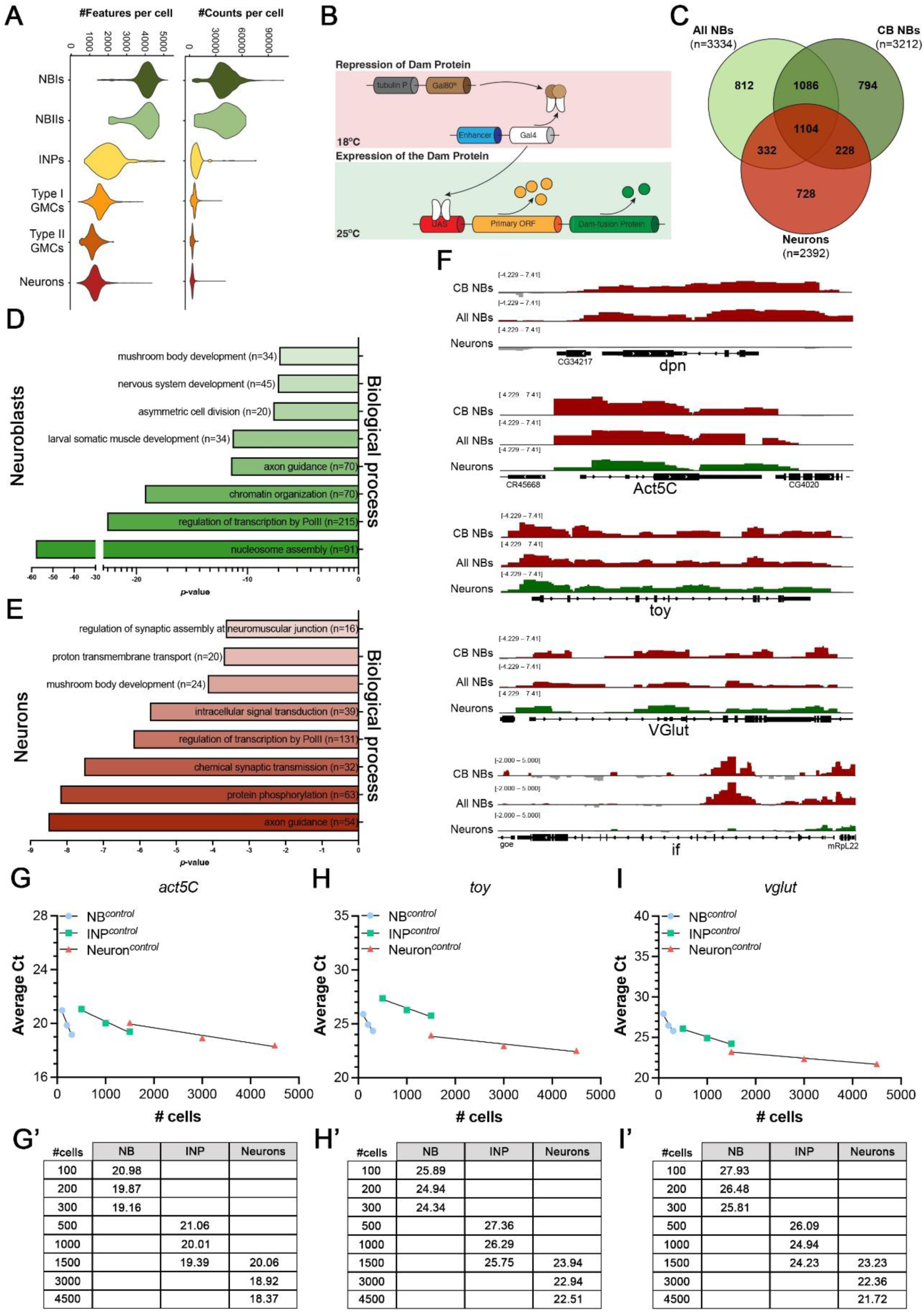
Neuroblasts exhibit higher transcriptional output and RNA Polymerase II recruitment compared to their differentiated progeny. **(A)** Single-cell RNA-sequencing (scRNAseq) analyses showing the number of features and counts per cell type in *D. melanogaster* neuroblast (NB) lineages. **(B)** Schematic representation of targeted DamID (TaDa) experimental design. **(C)** Venn diagram showing the number of genes bound by RNA Polymerase II (Pol II) in NBs and neurons, as determined by TaDa. Genes bound in NBs are shown in green: light green denotes all neuroblasts (All NBs, including both central brain and OL NBs, based on Southall *et al*.(Southall et al. 2013)); dark green indicates central brain neuroblasts (CB NBs) as determined in this study. Genes bound in neurons are shown in red, based on Southall *et al*.(Southall et al. 2013). **(D-E)** Gene ontology (GO) analysis of biological processes enriched among genes bound by Pol II in **(D)** NBs and **(E)** neurons. The *x*-axis shows log_10_(*p*-value) for each GO term. **(F)** Representative genome browser tracks of Pol II binding in ALL NBs, CB NBs, and neurons at selected *loci*: *dpn*, *act5C*, *toy*, *vglut,* and *inflate* (*if*). (**G-I’)** Quantification of gene expression by RT-PCR for (G–G’) *act5C*, (H–H’) *toy*, and (I–I’) *vglut* in FACS sorted NBs, intermediate neural progenitors (INPs), and neurons. Graphs show average Ct values (*y*-axis) plotted against the number of cells used per reaction (*x*-axis); corresponding Ct values are summarized in adjacent tables.

### Hypertranscriptional effects are not dependent on cell size

Recent studies suggest a positive correlation between cell size and transcription: cells with reduced transcription shrink to match lower activity, while increased transcription leads to larger cell volume, maintaining RNA concentration(Basier and Nurse 2023). Although this coupling has not been systematically and comparatively validated across diverse cell types, we asked whether the higher transcriptional output of NBs compared to neurons simply reflects their larger size. To separate transcriptional effects from size, we employed two complementary approaches. First, if transcription scales with cell size, closely related NBs of different sizes, like NBI and NBII, should differ in transcriptional output. Contrary to this expectation, scRNAseq analysis revealed that NBI and NBII display highly similar numbers of detected features and UMIs per cell (4016 *versus* 3834 features; 39427 *versus* 37205 UMIs), despite NBII being at least twice the volume of NBI(Homem et al. 2014) (Figure 1A and Supplemental Figure S1A). This demonstrates that hypertranscription in *Drosophila* neural stem cells can be uncoupled from cell size. Second, if cell size was a major determinant of transcriptional output, neurons of different sizes would be expected to show proportional differences in UMI counts per cell, even if the number of detected genes remained relatively constant. Using published scRNAseq data from adult *Drosophila* brains(Li et al. 2022) and volume estimates from publicly available electron microscopy datasets (https://codex.flywire.ai/) (Dorkenwald et al. 2024; Schlegel et al. 2024; Zheng et al. 2018), we analyzed transcription relative to neuronal subtype volume. Across different neuronal subtypes, both UMI counts and detected gene features exhibited highly similar distributions, irrespective of cell size (Supplemental Figure S1B–D), suggesting that transcriptional output is largely independent of neuronal volume. To further assess the relationship between cell size and transcriptional activity, we examined the association between average cell volume and mean UMI counts and detected features across subtypes (Supplemental Figure S1E). Overall, cell volume shows only a weak correlation with either the number of detected transcripts (*r* = 0.3425, *p*-value = 6.85 x 10^-4^) or the number of expressed genes (*r* = 0.2611, *p*-value = 6.92x 10^-3^). Notably, neuronal subtypes with markedly smaller volumes, such as T4/T5 from the optic lobe (OL), exhibit comparable, or higher numbers of UMIs and detected features than larger neurons such as L1 or L2 neurons (Supplemental Figure S1E, highlighted in red). Together, these analyses demonstrate that although NBs are substantially larger than neurons, their hypertranscriptional state cannot be explained simply by increased cell size, nor is transcriptional output directly proportional to cell volume across neuronal cell types.

### Drosophila neuroblasts exhibit RNA Polymerase II recruitment profiles consistent with hypertranscriptional states

Although scRNAseq provides an in-depth view of the transcriptional state of the cell, it has some caveats. Namely, changes in mRNA abundance do not always reflect alterations in transcriptional activity, as they may also result from post-transcriptional mechanisms affecting RNA stability. Hence, increased mRNA levels do not necessarily equate to increased transcription by Pol II. To further assess transcriptional activity in NBs and neurons, we used Targeted DamID (TaDa) (Figure 1B) to examine the genome-wide recruitment of Pol II in these cells. This method involves fusing a Pol II core subunit (RpII215) to the bacterial DNA adenine methyltransferase (Dam), which methylates adenines in GATC motifs. When Dam::Pol II binds to chromatin, it deposits methylation marks on nearby DNA, which can then be mapped by sequencing to infer Pol II occupancy.

To obtain NB-specific profiles of Pol II, we expressed the Dam-fused PolII under the control of a CB NB-specific GAL4 (VT201094), the same driver used to generate the scRNAseq dataset(Marques et al. 2023). Additionally, we analyzed previously published Pol II TaDa datasets that include both OL and central-brain (CB) NBs (using *worniu-GAL4*, indicated in the Figure as All NBs) and another dataset that includes neurons (using the pan-neuronal driver *elav-GAL4*)(Southall et al. 2013). As expected, and while sharing approximately 70% of common genes bound by Pol II, the TaDa profiles obtained using different NB-specific Gal4 drivers (*worniu-GAL4 versus VT201094-GAL4*) rendered distinct binding signatures, reflecting the differences in the NB populations analyzed (Figure 1C). Consistent with the scRNAseq data, the number of Pol II-bound genes was much higher in NBs in comparison to neurons (3224 and 3212 *versus* 2392 genes, respectively; Figure 1C). These findings demonstrate that NBs exhibit both greater Pol II occupancy on chromatin and increased number of expressed genes, supporting the conclusion that *Drosophila* NSCs exist in a hypertranscriptional state.

A key feature of hypertranscription is the expression of genes that are not functionally required for a particular cell state, such as the expression in NSCs of genes specific for differentiated neurons, muscle or germline. To determine whether such pattern of gene expression occurs in *Drosophila* NBs, we performed gene ontology (GO) enrichment analysis to identify biological processes overrepresented in the PolII TaDa datasets from CB NBs and neurons (Figure 1D and 1E). As expected, genes involved in asymmetric cell division and nervous system development were significantly enriched in NBs (Figure 1D). This enrichment is validated by genome-browser tracks showing robust Pol II recruitment to canonical NB-expressed *loci*, such *deadpan* (*dpn*)(Figure 1F, first panel)(Homem et al. 2015). Pol II binding is detected at *dpn locus* in NBs but is absent in neurons. Interestingly, genes typically associated with other cell types, such as those involved in muscle development or neuronal function, were also overrepresented among NB-bound Pol II targets, suggesting that Pol II is recruited to unexpected genomic regions in these cells (Figure 1D). For example, *twin of eyeless* (*toy*), the *Drosophila* orthologue of PAX6, plays a key role in retinal determination and central nervous system development and is typically restricted to neurons(Czerny et al. 1999). Yet *toy* shows Pol II occupancy not only in neurons, but also in NBs (Figure 1F). Similarly, *Vglut*, a vesicular glutamate transporter specifically expressed in glutamatergic neurons(Deshpande et al. 2020), also exhibits Pol II binding in NBs as well as neurons (Figure 1F, third and fourth panel). Another illustrative example is *inflate* (*if*), which encodes αPS2, a component of the αPS2βPS integrin required for muscle attachment, midgut morphogenesis, and epithelial adhesion between wing epithelia in the adult fly(Devenport et al. 2007). Pol II is robustly recruited to *if* in NBs but not in neurons (Figure 1F, bottom panel). These observations further support the conclusion that NBs have a broad, non-cell type specific transcriptional program.

Hypertranscription has also been associated with enhanced expression of genes involved in chromatin remodeling and epigenetic plasticity(Percharde et al. 2017). Consistent with this, GO terms related to nucleosome assembly and chromatin organization were also enriched among Pol II-bound genes in NBs, but not in neurons (Figure 1D and 1E). In neurons, as expected, Pol II-bound genes were enriched for functions related to axon guidance, chemical synaptic transmission, and synaptic assembly (Figure 1E).

Altogether, our scRNAseq and TaDa analyses demonstrate that *Drosophila* NBs exhibit hallmark features of hypertranscription: they express more genes and transcripts than their differentiated progeny, show elevated genome-wide Pol II recruitment, and display widespread Pol II binding to genes beyond their expected lineage, including neuron- and muscle-specific *loci*, revealing a broad, non–cell type–restricted transcriptional program in NSCs.

### Detection of hypertranscription in NBs using quantitative gene expression analysis

Normalization methods commonly used in transcriptomic studies, such as referencing housekeeping genes or total RNA content, often mask hypertranscriptional states, making them difficult to detect and study(Kim et al. 2024, 2023). To overcome this limitation, more precise approaches like scRNAseq or quantitative real-time PCR (RT-PCR) using exact cell numbers offer greater accuracy for absolute quantification.

To assess expression differences between NBs and differentiating/differentiated progeny, we performed RT-PCR using defined number of sorted cells (Figure 1G-I’). Specifically, we collected brains from wandering third instar larvae expressing membrane-localized GFP under the control of the CB NBs-specific driver (*VT201094*-*GAL4*), which were chemically and mechanically dissociated. Since the GFP expressed in the NBs is inherited by their progeny, this effectively labels their lineage including neurons. These cells were collected via FACS based on size and fluorescence: NBs are the largest and brightest, INPs are intermediate, and neurons are the smallest and dimmest(Harzer et al. 2013). RNA was extracted from sorted cells and used for RT-PCR. The number of cells used per RT-PCR reaction was determined based on both empirical and theoretical considerations, ensuring quantification within the linear range of the assay. Due to the relative scarcity of NBs, we used 100, 200, or 300 cells per reaction; for INPs, 500, 1000, or 1500 cells; and for neurons, which are more abundant, 1500, 3000, or 4500 cells.

To determine expression levels in NBs and in their differentiated progeny, we examined the expression of three genes: the housekeeping gene, *act5C* and two neuron-associated genes, *toy* and *vglut*. *Act5C* was chosen as a housekeeping control for general transcriptional activity. *Toy* and *vglut* were selected because, although they are typically expressed in differentiated neurons, they are expressed in NBs under hypertranscriptional conditions. The expression levels of genes not normally required in progenitor cells may serve as indicators of defects in hypertranscription regulation. To compare the expression levels of each gene among the different cell types, two strategies were used: (i) comparing average C_t_ values using the same numbers of cells (*e.g.* C_t_ for 1500 INPs *vs.* 1500 neurons); (ii) determining the number of cells required to obtain similar C_t_ values (*e.g.* similar C_t_ for 300 NBs and 3000 neurons). As expected, *act5C* is expressed in all cell types (Figure 1G), however, its expression levels varied significantly. For instance, 300 NBs had a Ct of 19.16, similar to 3000 neurons with a Ct of 18.92, indicating approximately 10-fold higher expression in NBs. Similarly, *act5C* expression in 100 NBs (Ct = 20.98) matched that of 500 INPs (Ct = 21.06), reflecting a 5-fold higher expression in NBs. Lastly, *act5C* expression in INPs was 1.6-fold higher than in neurons, as observed when comparing 1500 INPs and 1500 neurons (2^-ΔCt^ = 2^-(Ct^ ^INPs-Ct^ ^neurons)^ = 2^-(19.39-20.06)^ = 1.59) (Figure 1G’). These results indicate a progressive decrease in *act5C* expression from NBs to INPs to neurons, consistent with a decline in transcriptional activity during differentiation.

*toy* was also found to be expressed in NBs, INPs and neurons (Figure 1H). *toy* expression was also higher in NBs when compared to INPs: 100 NBs (C_t_=25.89) matched 1500 INPs (C_t_=25.75), indicating a ∼15-fold difference. In neurons, *toy* was more expressed than in INPs (neurons C_t_=23.94 *vs.* INP C_t_=25.75 for 1500 cells), a 3.5-fold reduction in INPs *vs.* neurons (2^-ΔCt^ = 2^-(Ct^ ^neurons-Ct^ ^INPs)^ = 2^-(23.94-25.75)^ = 3.51) (Figure 1H’). While no direct comparison between NBs and neurons is possible to be made, given that we know how NBs and neurons compare to INPs, we can infer a comparison between these two cell types: *toy* is approximately 4.3 times more expressed in NBs than in neurons. This exacerbated expression of *toy* in NBs is likely due to the underrepresentation of toy-positive neurons among the purified neuronal population, since this gene is not expressed in all neuronal subtypes or developmental times(Furukubo-Ttokunaga et al. 2009). *vglut* transcripts were also detected in NBs, INPs, and neurons (Figure 1I). *vglut* expression was 2-fold higher in neurons than INPs (C_t_=23.23 *vs.* C_t_=24.23 for 1500 cells, 2^-ΔCt^ = 2^-(Ct neurons-Ct INPs-)^ = 2^-(23.23-24.23)^ = 2). Moreover, *vglut* expression was approximately 2-fold higher in NBs compared to INPs (Ct = 26.48 and 25.81 for 200 and 300 NBs, respectively, *vs.* 26.09 for 500 INPs) (Figure 1I’). Interestingly, *vglut* expression levels were similar between NBs and neurons, inferred from their similar ratio of *vglut* expression *vs.* INPs. This could be explained by the fact that, when sorting neurons by FACS, we collect a mixture of all neurons and not only *vglut*-expressing ones (glutamatergic neurons), which may dilute *vglut* expression(Lacin et al. 2019).

Additionally, since different numbers of cells were used per RT-PCR reaction, we were able to fit our results to a mathematical approximation. This allowed us to assess whether our empirical observations aligned with the mathematical model. For *act5C*, curve fitting revealed that this gene is expressed 5 times more in NBs compared to INPs, and 1.6 times more in INPs than in neurons (Supplemental Figure S2A–A’; compare empirical results – red – with the mathematical calculation – green). Consistent with our previous findings, *toy* is more highly expressed in NBs and neurons than in INPs (10-fold and 3.5-fold, respectively) (Supplemental Figure S2B–B’). Finally, regarding *vglut* expression, INPs show lower levels compared to both NBs and neurons (Supplemental Figure S2C–C’). Altogether, these results demonstrate that hypertranscription occurs in NBs, characterized by higher number of genes expressed, elevated expression of housekeeping gene, transcription of genes that are not typically expected to be expressed in NBs and broader genome-wide Pol II binding. This transcriptional intensity appears to diminish as cells differentiate, supporting a model where hypertranscription is associated with cell potency.

### The Mediator complex is a regulator of hypertranscription in neuroblasts

The elevated transcriptional output observed in NBs, both at the level of gene expression and Pol II recruitment, suggests the presence of a specialized transcriptional amplification program that supports SC identity and function. However, the molecular mechanisms driving this hypertranscriptional state remain largely unclear. Given Mediator’s central role in regulating distinct transcriptional states in SCs and their differentiated progeny, as well as its established function in differentiation, we hypothesized that this coactivator complex is essential for establishing hypertranscriptional programs in SCs.

To investigate whether Mediator is a regulator of hypertranscription we examined Mediator’s chromatin binding profile in larval CB NBs using TaDa (Dam-Mediator subunit driven by *VT201094-GAL4*). Given the modular structure of the Mediator complex, comprising head, middle, tail, and kinase modules(Asturias et al. 1999), we selected and generated Dam fusions for representative subunits from distinct modules (Med11 and Med17 from the head, Med21 from the middle, and Med27 from the tail) to gain comprehensive insight into its genome-wide occupancy. Each subunit displayed a distinct but overlapping binding profile, with thousands of gene targets: 2235 for Med11, 2016 for Med17, 2136 for Med21, and 2503 for Med27 (Figure 2A). Approximately 50% of genes were co-bound by all subunits, while several hundred exhibited subunit-specific binding. Given that Mediator is a large 1-2.5MDa complex with varying subunit accessibility to DNA, these binding profiles likely reflect differences in chromatin accessibility rather than distinct functional interactions(Robinson et al. 2016). Therefore, we considered that the overall binding pattern of Mediator is best represented by the combined set of peaks identified across all subunits. Taking that into consideration, we have identified 3714 unique genes bound by Mediator, confirming its genome-wide regulatory role in NBs. GO analysis revealed enrichment for key neurodevelopmental biological processes, including asymmetric cell division (Figure 2B), cell differentiation (Figure 2C and 2E), and central and peripheral nervous systems development (Figure 2B and 2D). Additionally, there was also strong enrichment for genes associated with regulation/positive regulation of transcription by Pol II, aligning with the hypothesis that Mediator contributes to hypertranscription in NBs. Like Pol II, Mediator also bound unexpected genes, including those typically linked to differentiated cell types, such as genes involved in axon guidance, neuron differentiation, dendrite morphogenesis, and the regulation of neurogenesis (Figure 2B-E). Notably, just like Pol II, all tested Mediator subunits showed robust binding not only to *act5C* but also to unexpected targets such as *toy* and *vglut* (Figure 2F), despite these genes being functionally unrelated to NB self-renewal. Collectively, these results demonstrate that Mediator exhibits a broad chromatin binding profile, including binding to unexpected genes, similar to Pol II in NBs, supporting its potential role in regulating hypertranscription.

**Figure 2.**
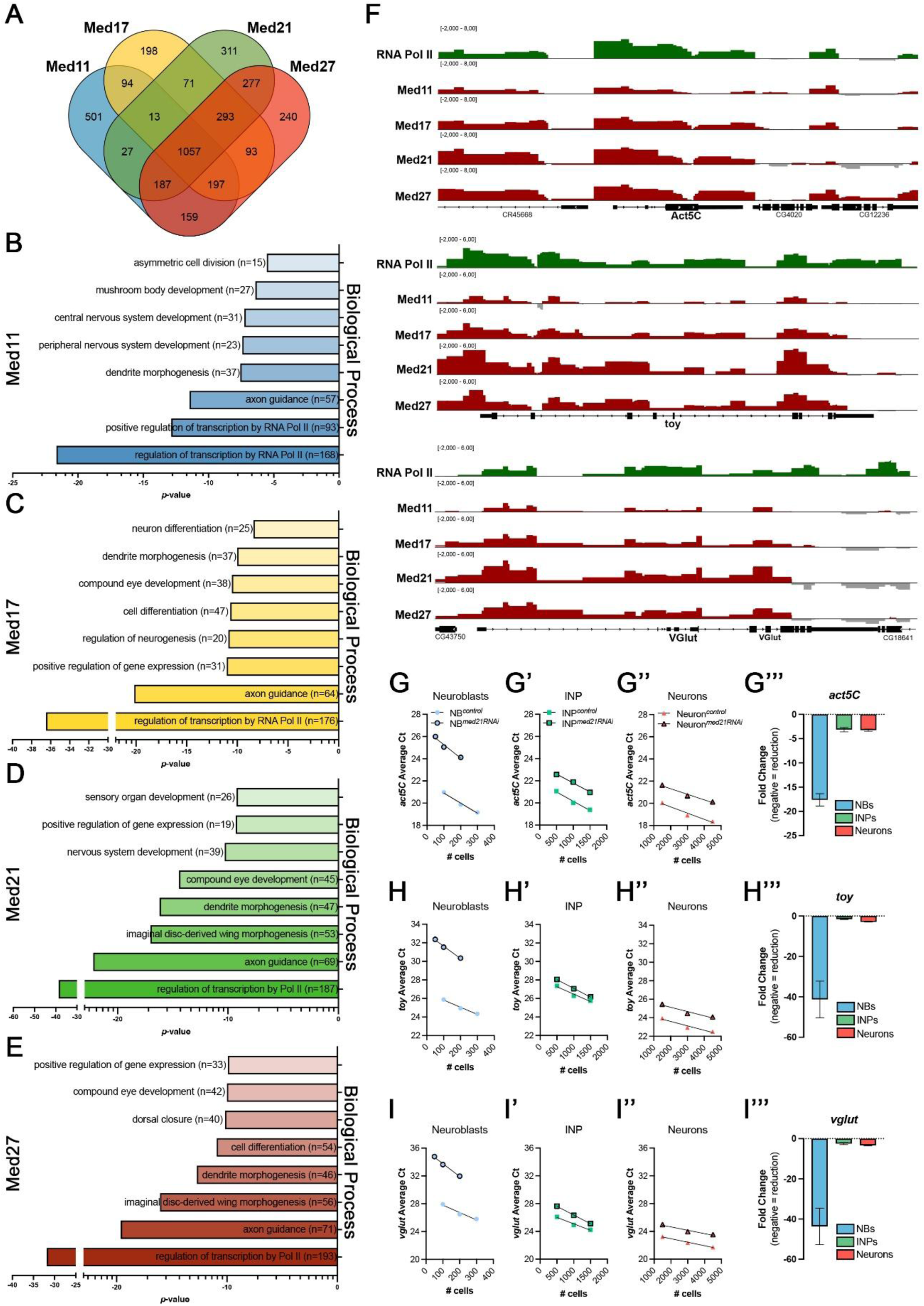
The Mediator complex is a pivotal regulator of hypertranscription in neuroblasts. **(A)** Venn diagram showing the number of genes bound by individual subunits Med11 (blue), Med17 (yellow), Med21 (green), and Med27 (red) in central brain neuroblasts (NBs), as determined by TaDa. Data representative of two biologically independent experiments. **(B-E)** Gene ontology (GO) analysis of biological processes significantly enriched among genes bound by **(B)** Med11, **(C)** Med17, **(D)** Med21, and **(E)** Med27. *x*-axis represents the log_10_(*p*-value) for each term. **(F)** Representative genome browser tracks showing Polymerase II (green) and Mediator subunits (red) occupancy at the *act5C*, *toy*, and *vglut loci* in NBs. **(G-G’’’)** Quantification by RT-PCR of *act5C* expression in FACS sorted **(G)** neuroblasts (NBs), (**G’)** intermediate neuronal precursors (INPs), and **(G’’)** neurons in control NBs and upon *med21* RNAi. **(G’’’)** Average fold decrease values (mean ± SD) for *act5C* expression in each cell type upon *med21* depletion (in comparison to control, no change = 0). **(H-H’’’)** Quantification of *toy* expression in **(H)** NBs, (**H’)** INPs, and **(H’’)** neurons upon RNAi of *med21*. **(H’’’)** Average fold decrease values (mean ± SD) for *toy* expression in each cell type upon *med21* depletion (in comparison to control, no change = 0). **(I-I’’’)** Quantification of *vglut* expression in **(I)** NBs, (**I’)** INPs, and **(I’’)** neurons upon *med21* RNAi. **(I’’’)** Average fold decrease values (mean ± SD) for *vglut* expression in each cell type upon *med21* depletion (in comparison to control, no change = 0). RT-qPCR data are plotted with the number of cells used per reaction on the *x*-axis and the corresponding average Ct value on the *y*-axis. Fold decrease was determined using the ΔCt method.

Next, we functionally tested whether Mediator affects hypertranscription in NBs. To do this, we reduced Mediator levels by RNAi and compared transcriptional levels between NBs and more differentiated progeny. Specifically, we targeted the Med21 subunit, identified in the TaDa analyses as a key component of the Mediator complex. Using a CB NB-specific driver (*VT201094-GAL4*) we express UAS-RNAi to deplete Mediator activity in the NB lineage. Although the driver is specific to NBs, GAL4 and GFP are inherited by their progeny, ensuring efficient knockdown across the entire lineage (Supplemental Figure S3). To assess the impact of Mediator depletion on transcriptional activity, we sorted NBs, INPs, and neurons by FACS and performed RT-qPCR on an exact number of cells to quantify the expression of *act5C, toy*, and *vglut* (Figure 2G-I’’’). Loss of Med21 resulted in a dramatic reduction in gene expression in NBs, with *act5C* downregulated 17.6-fold, *toy* by 41.3-fold, and *vglut* by 43.7-fold (Figure 2G and G’’’, H and H’’’, I and I’’’). In contrast, expression changes in INPs and neurons were much more modest, ranging from 1.54- to 3.30-fold) (Figure 2G’-I’’’).

These results indicate that Mediator plays an essential role in sustaining high levels of transcriptional activity specifically in NBs, with a notably smaller impact on differentiating or differentiated cells. Together, these findings establish Mediator as a central regulator of hypertranscription in *D. melanogaster* NSCs.

### Mediator-dependent hypertranscription is required for proper neuroblast division and progeny differentiation

The functional significance of hypertranscription *in vivo*, particularly its effects cell identity and differentiation, remains poorly understood(Kim et al. 2024). Our findings showed that NBs are in a hypertranscriptional state, supported by broad Mediator recruitment. To investigate the *in vivo* role of hypertranscription, we selectively depleted core Mediator subunits from both type II and type I NBs.

First, we systematically depleted individual Mediator subunits in type II NBs using the NBII-specific driver *worniu-GAL4, asense-GAL80*, and assessed lineage composition with cell-type specific markers (Figure 3A). For most subunits tested, depletion did not result in any obvious phenotype compared to controls, with lineages maintaining a single Dpn⁺/Ase⁻ NB and the expected composition of intermediate progenitors (Supplemental Figure S4A). However, depletion of four subunits – *Med11, Med14, Med17,* and *Med21* – resulted in smaller lineages with multiple Dpn⁺/Ase⁻ NB-like cells (white arrows), accumulation of immature Ase⁻ INPs (orange arrows), and a concomitant loss of Ase⁺ INPs (yellow arrows) and mature Ase+Dpn+ INPs (pink arrows) (Figure 3B, Supplemental Figure S4B, quantified in Figure 3C–F).

**Figure 3.**
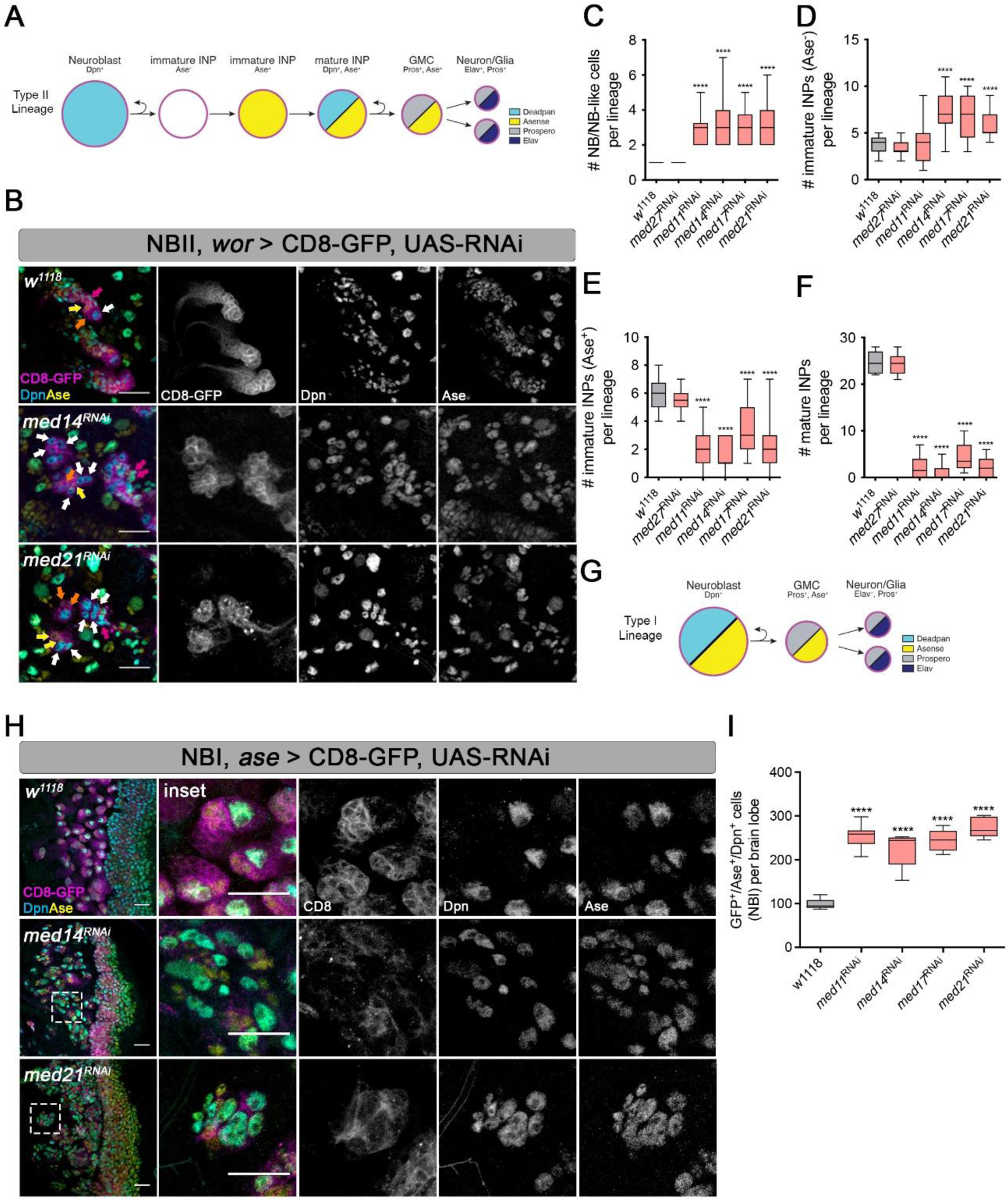
Mediator depletion results in abnormal neuroblasts lineage progression. **(A)** Schematic representation of type II neuroblast (NB) lineages. Cell types are color-coded according to specific markers used for their identification. The magenta outline indicates GFP expression, marking the cells where the driver and RNAi constructs are expressed. **(B)** Immunostaining of fixed wandering third instar (L3) brains expressing UAS-CD8::GFP (magenta) and RNAi against *med14* (middle panel) or *med21* (bottom panel) under the control of the type II-specific driver *worniu-GAL4 asense-GAL80*. Control brains (top panel) is *w^1118^*. Cell identity was assessed using Deadpan (Dpn, cyan) and Asense (Ase, yellow) immunostaining. White arrows indicate Dpn-positive NBs; orange arrows, Ase-negative immature INPs; yellow arrows, Ase-positive immature INPs; pink arrows mature INPs co-expressing Dpn and Ase. Scale bar = 20μm. **(C-F)** Quantification of the number of **(C)** NBs, **(D)** Ase-negative immature INPs, **(E)** Ase-positive immature INPs, **(F)** mature INPs upon depletion of *med27*, *med11*, *med14*, *med17*, and *med21* from type II NBs. Box-plot representing data distribution, with box being representative of the interquartile range containing the middle 50% of the dataset and mid-line representing the median value. Lines extending from the box are indicative of the range of the dataset. n ≥ 5 and at least 4 independent lineages per one lobe were analyzed. Statistical significance calculated using one-way ANOVA. **** = *p* < 0.0001. **(G)** Schematic representation of type I NB lineages. Cells are colored according to the specific markers that allow for their accurate identification. The magenta outline indicates GFP expression, marking the cells where the driver and RNAi constructs are expressed. **(H)** Immunostaining of L3 larval brains expressing UAS-CD8::GFP (magenta) and RNAi against *med14* (middle) and *med21* (bottom) under the control of the type I-specific driver (*asense-GAL4*). Top panel represents the control (*w^1118^*). Dpn (cyan) and Ase (yellow) staining were used to identify type I NBs. Dashed boxes indicate the regions shown in the insets. Scale bar = 20μm. **(I)** Quantification of the total number of Ase- and Dpn-positive type I NBs upon *med11*, *med14*, *med17*, and *med21* RNAi. Box-plot representing data distribution, with box being representative of the interquartile range containing the middle 50% of the dataset and mid-line representing the median value. Lines extending from the box are indicative of the range of the dataset. n = 8 for all tested conditions. Statistical analysis was performed using one-way ANOVA. **** = *p* < 0.0001.

These defects were recapitulated in type I lineages upon Mediator depletion using the *asense-GAL4* driver. While the depletion of the majority of Mediator subunits did not cause a phenotype (Supplemental Figure S5A), knock down of *Med11*, *Med14*, *Med17* and *Med21* in type I lineages also led to lineages containing several ectopic NBI-like cells (Figure 3G, H; Supplemental Figure S5B), and a twofold increase in the total number of NBs per lobe (Figure 3I). Additionally, brain lobes were overall smaller following Mediator depletion in NBIs, suggesting a failure in producing differentiated neurons in normal numbers.

Previously we have shown that Mediator-dependent hypertranscription is exclusive to NBs but not INPs and neurons. Indeed, depletion of Mediator from NBs leads to dramatic lineage defects. However, since *NB-GAL4* drivers also target INPs and neurons (Supplemental Figure S3), it is possible that the observed phenotypes might result from Mediator depletion in these differentiating cells rather than in NBs themselves. To test this, we specifically depleted Mediator subunits in INPs and neurons using cell type-specific drivers. The *earmuff-GAL4* driver, which specifically targets immature INPs (Ase⁻ or Ase⁺, depending on the insertion), and the *nsyb-GAL4* driver, which targets neurons, were used to drive RNAi against Mediator subunits. In both cases, knockdown of *Med27*, *Med11*, *Med14*, *Med17*, or *Med21* did not result in any detectable phenotype (Supplemental Figure S6A–C). Together, these results confirm that the differentiation defects are specifically due to Mediator depletion in NBs, rather than in downstream progenitor cells, underscoring the crucial role of Mediator specifically in NBs for lineage differentiation.

Interestingly, while Mediator depletion in NBs results in the formation of multiple NB-like cells, it does not lead to tumor overgrowth, as seen in classical NB fate mutants such as *brat*, *miranda*, or *numb*(Homem and Knoblich 2012). This suggests that these NB-like cells are either eliminated by apoptosis or fail to proliferate efficiently. To assess apoptosis, we stained brains Mediator-depleted brains for Death caspase 1 (Dcp-1), a key effector caspase involved in protein cleavage(Song et al. 1997). While Dcp-1 staining reliably labels apoptotic neurons under normal conditions, no Dcp-1 positive NBs were observed upon depletion of *med14* or *med21* (Figure 4A). Thus, we can rule out apoptosis as the cause of the absence of tumor formation in Mediator-depleted NBs. We next tested whether NB proliferation was affected by staining for phospho-histone H3 (pH3), a mitotic marker(Hendzel et al. 1997) (Figure 4B). Given the increased number of NB-like cells upon Mediator depletion, we quantified the mitotic rate as the ratio of pH3-positive to the total number of NB/NB-like cells. Knockdown of *med14* or *med21* resulted in a significant decrease in the mitotic rate, which dropped to 13.32% and 12.16%, respectively, compared to 25% in controls (Figure 4B). This indicates that Mediator-depleted NBs divide more slowly, at least partially explaining the lack of tumor formation despite the increased number of NB-like cells.

**Figure 4.**
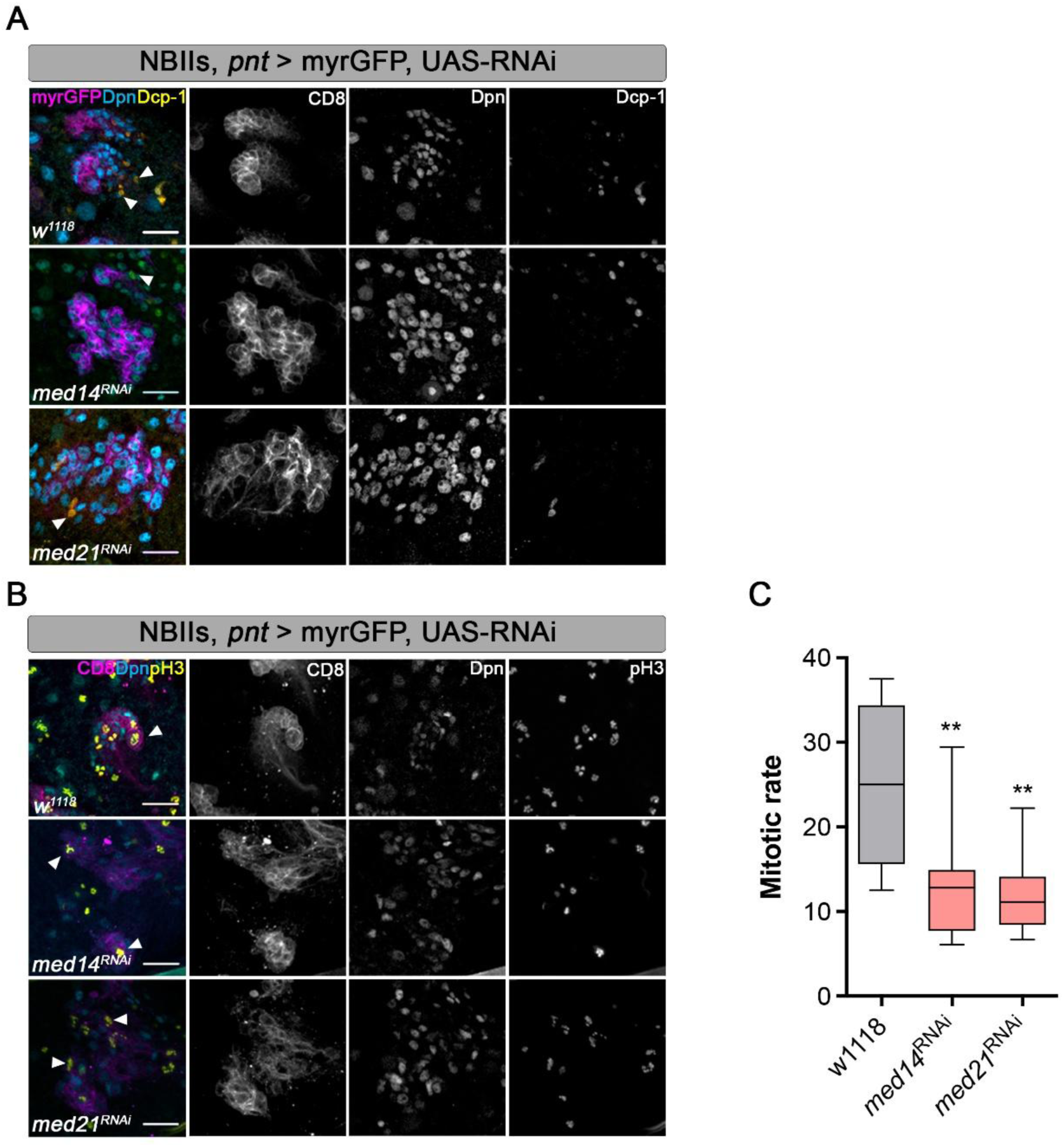
Mediator depletion reduces the mitotic activity of neuroblasts. **(a-b)** Confocal images of fixed wandering 3^rd^ instar (L3) brains expressing UAS-myristoylated::GFP (myrGFP, magenta) and RNAi against *med14* (middle panel) or *med21* (bottom panel) under the control of the type II-specific driver *pointed-GAL4*. *w^1118^*was used as a control (top panel). Scale bars = 20μm. **(A)** Neuroblasts (NBs) were identified by deadpan staining (Dpn, cyan), and apoptotic cells were labeled with Dcp-1 (yellow). **(B)** Brains stained with Dpn to identify NBs (Dpn, cyan) and phosphorylated histone H3 (pH3, yellow) to label actively dividing cells. **(C)** Quantification of NB mitotic rate upon RNAi of *med14* or *med21*. Mitotic rate was calculated as the percentage of pH3-positive NBs among the total number of type II NBs. n ≥ 5 and statistical significance calculated using one-way ANOVA. ** = *p* < 0.01.

Collectively, our results suggest that Mediator-dependent hypertranscription in NBs is essential for sustaining the transcriptional program that enables both their proliferation and proper differentiation. Loss of Mediator disrupts this balance, resulting in slow-dividing, supernumerary NBs, with impaired lineage progression. Notably, although NB-like cells accumulate, the associated proliferation defects likely prevent tumor overgrowth, highlighting Mediator’s dual role in promoting NB expansion while safeguarding timely differentiation.

### Metabolic perturbation does not impair NB hypertranscription

Stem cell function depends on the tight coordination of multiple cellular processes, including transcription and metabolism. It has been proposed that substantial energetic demand associated with hypertranscription requires the precise coordination of metabolic activity to sustain the cellular processes underlying high transcriptional output(Bulut-Karslioglu et al. 2018b; Kim et al. 2024). In *Drosophila*, Mediator has been shown to regulate both transcription and metabolism in pupal NBs, promoting a switch from glycolysis to oxidative phosphorylation (OxPhos) to support NB terminal differentiation(Homem et al. 2014). Given Mediator’s dual roles, we hypothesized that Mediator might support NB identity and differentiation by coordinating metabolic activity with transcriptional regulation.

Consistent with this idea, chromatin profiling revealed that multiple Mediator subunits bind genes encoding key enzymes of glycolysis and the TCA cycle, such as *ald1*, *gapdh1*, *hex-A*, and *pyk*, suggesting potential direct regulation of metabolic pathways (Figure 5A). Moreover, since metabolites such as ATP and acetyl-CoA can influence chromatin state and gene expression, we wondered whether metabolism could feed back to support hypertranscription in NBs. To determine whether Mediator loss alters metabolic activity, we assessed whole-brain metabolism following NB-specific knockdown of representative Mediator subunits. Seahorse analysis showed a shift toward glycolysis (increased ECAR and reduced OCR) upon *med17* or *med21* depletion (Supplemental Figure S7A–C). While this shift may partly reflect changes in lineage composition – namely, a reduction in OxPhos-reliant differentiated cells – we also detected metabolic changes at the NB level using genetically encoded biosensors. In NBII lineages depleted for *med21*, intracellular glucose levels were reduced, consistent with either decreased glucose uptake or increased consumption. However, lactate levels remained unchanged, suggesting that glucose was not diverted into anaerobic glycolysis, but potentially into biosynthetic pathways such as the pentose phosphate pathway (Supplemental Figure S7D–E).

**Figure 5.**
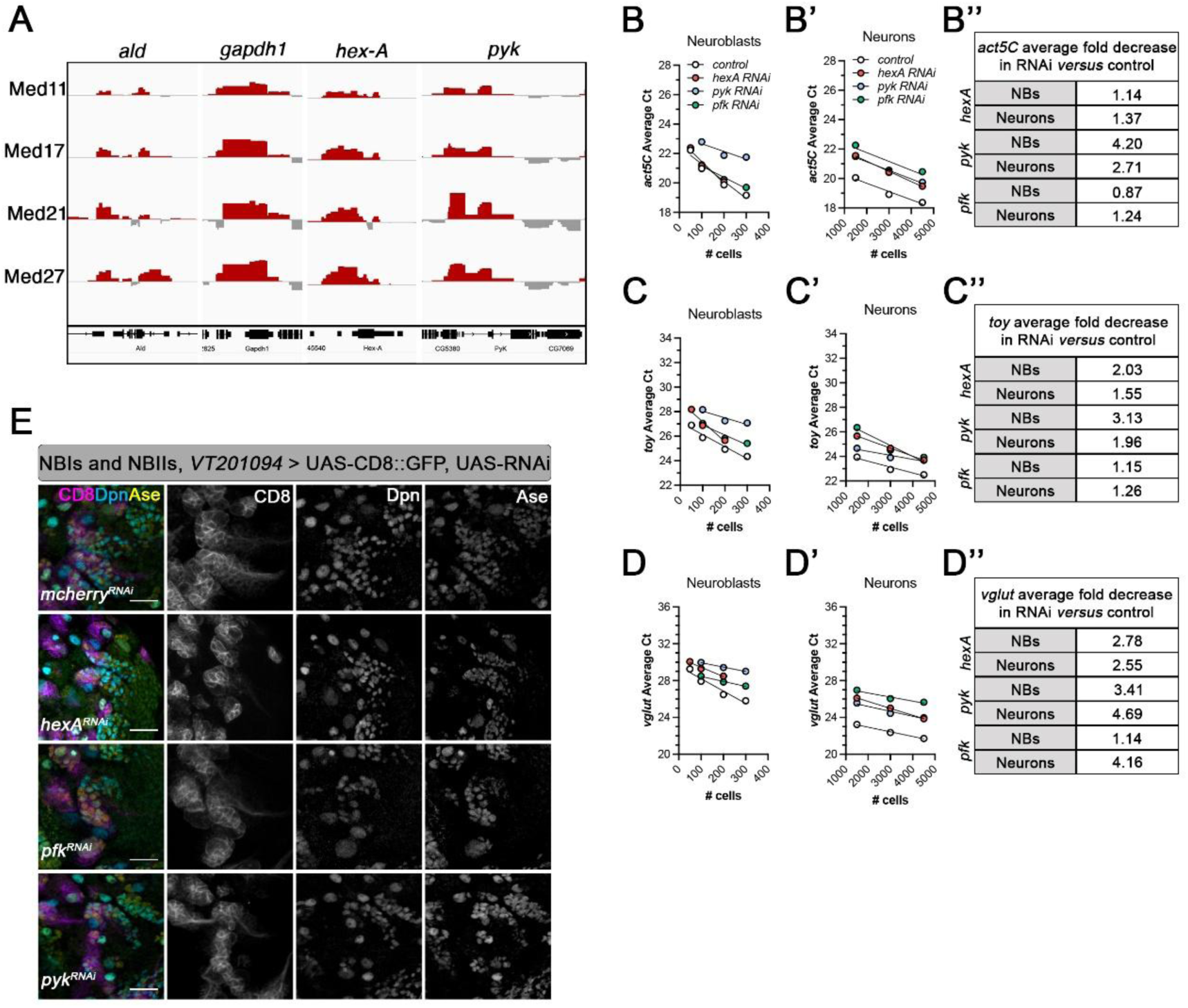
Metabolism alone is not sufficient to sustain hypertranscription. **(A)** Representative genome browser tracks showing recruitment of Mediator subunits to *loci* encoding metabolic enzymes, including *ald*, *gapdh1*, *hex-A*, and *pyk*. **(B-B’’)** Quantification by RT-PCR of *act5C* expression in FACS sorted **(B)** neuroblasts (NBs) and **(B’)** neurons upon RNAi-mediated depletion of *hex-A*, *pyk*, or *pfk* driven by *VT201094-GAL4*. **(B’’)** Corresponding table of average fold decrease between the RNAi against metabolic enzymes and control. **(C-C’’)** Quantification of *toy* expression by RT-PCR in FACS sorted **(C)** neuroblasts (NBs) and **(C’)** neurons upon RNAi-mediated depletion of *hex-A*, *pyk*, or *pfk* driven by *VT201094-GAL4.* **(C’’)** Corresponding table of average fold decrease between the RNAi against metabolic enzymes and control. **(D-D’’)** Quantification of *vGlut* expression by RT-PCR in FACS sorted **(D)** neuroblasts (NBs) and **(D’)** neurons upon depletion of *hex-A*, *pyk*, or *pfk* driven by *VT201094-GAL4*. **(D’’)** Corresponding table of average fold decrease between the RNAi against metabolic enzymes and control. **(E)** Confocal images of fixed wandering third instar (L3) brains expressing UAS-myr::GFP (magenta) and RNAi against *hex-A*, *pfk*, and *pyk* driven by CB NBs-specific *VT201094-GAL4*. Top panel shows control (*mcherry*RNAi). Staining with Deadpan (Dpn - cyan) and Asense (Ase - yellow) was used for proper identification of cell identity. Scale bar = 20μm.

To directly test whether metabolic activity is required to sustain hypertranscription, we perturbed glycolysis by targeting key glycolytic enzymes (*hex-A*, *pyk*, and *pfk*) using RNAi driven by a NB-specific driver (*VT201094-GAL4*). As hexokinase (*hex-A*) catalyzes the first step of glycolysis, phosphorylating glucose to glucose-6-phosphate, its knockdown is expected to block glucose utilization. Using a fluorescent glucose biosensor(Gándara et al. 2019a), we confirmed that *hex-A* depletion led to glucose accumulation in NBII lineages, indicating effective inhibition of glycolytic flux (Supplemental Figure S7F). We then tested whether reduced glycolysis affects hypertranscription. NBs and neurons were separately sorted following knockdown of each enzyme, and transcriptional output was assessed by RT-qPCR for selected genes. Surprisingly, gene expression was only mildly affected, with reductions of up to 4.2-fold in NBs and 2.7-fold in neurons (Figure 5B–D’’). In addition, NB lineage architecture remained intact, with no major defects observed in either NBI or NBII lineages (Figure 5E). Together, these results demonstrate that although Mediator regulates NB metabolism, hypertranscription is remarkably resilient to acute metabolic perturbations. This suggests that the mechanisms sustaining high transcriptional output in NBs are largely independent of metabolic state, at least within the physiological range tested.

### Mediator-dependent hypertranscription is required for growth in Drosophila brain tumors

Hypertranscription has recently emerged as a defining feature of aggressive human cancers, profoundly influencing tumor progression and clinical outcomes(Zatzman et al. 2022; Cao et al. 2022). Elevated transcriptional output is observed in approximately 80% of cancers – including breast, colorectal, uterine, and head and neck cancers(Zatzman et al. 2022). Given our findings that Mediator promotes hypertranscription in NSCs, we hypothesized that *Drosophila* NB-derived brain tumors similarly rely on Mediator-dependent hypertranscription for growth. In *Drosophila*, NSC-derived brain tumors can be readily induced by knocking down the tumor suppressor gene *brain tumor* (*brat*) in NBII cells(Homem and Knoblich 2012). To test whether tumor NBs exhibit elevated transcriptional activity compared to wildtype NBs, we performed scRNAseq on brains from control and *brat*-RNAi animals (Supplemental Table S1). Tumor-derived NBs (tNBs) displayed a significantly higher number of detected genes (median of 2811.5 vs. 2009) and increased total transcript counts per cell (median of 22,441 vs. 11,003) compared to wild-type NBs (Figure 6A–B), indicating that tNBs exist in an enhanced hypertranscriptional state.

**Figure 6.**
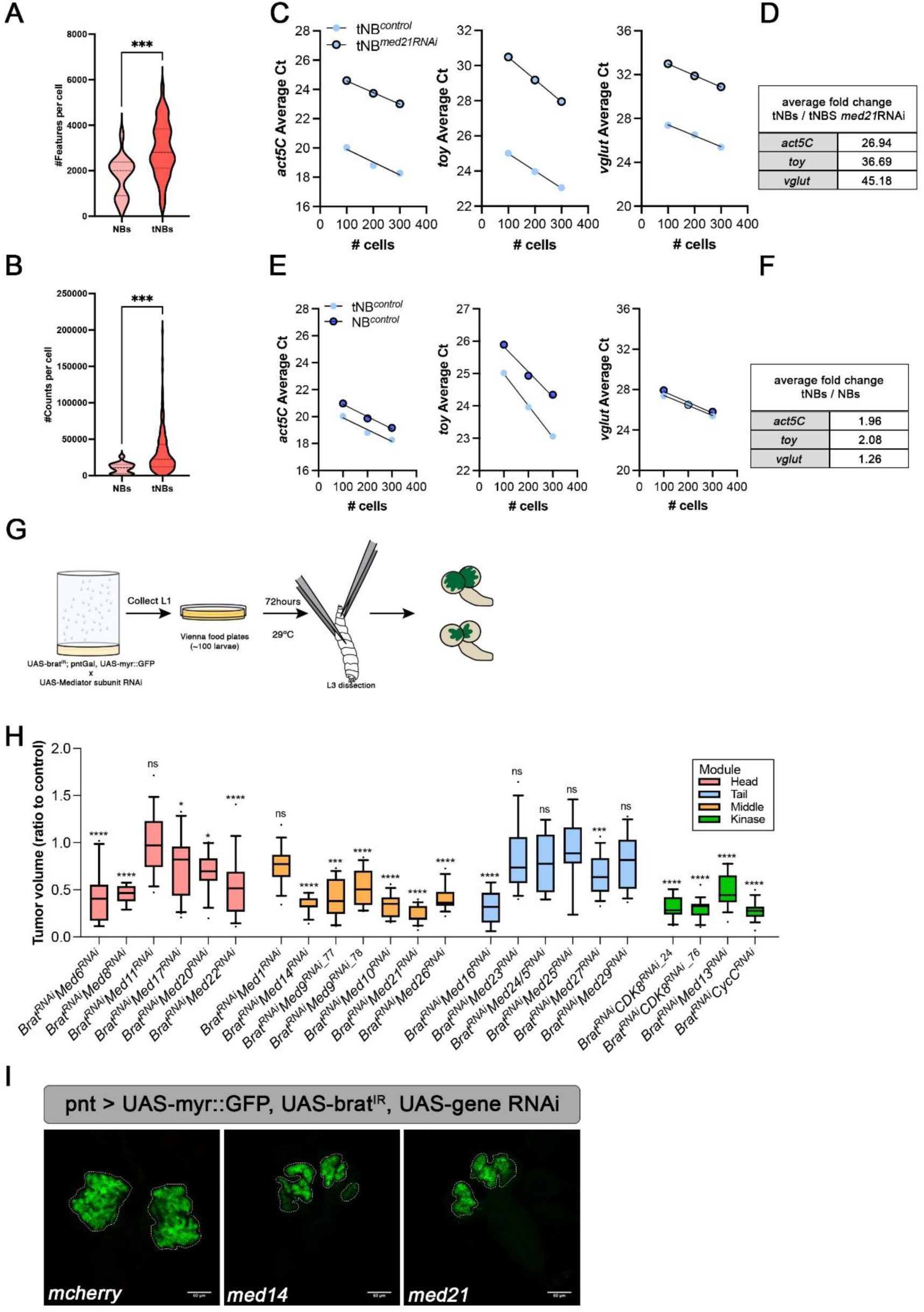
Hypertranscription in tumor-derived neuroblasts is dependent on the Mediator complex. **(A-B)** Violin plots displaying the **(A)** number of features and **(B)** number of counts per cell for wild-type neuroblasts (NBs, light coral) versus tumor-derived neuroblasts (tNBs, dark coral). Statistical significance was determined using an unpaired two-tailed *t* test. *** = *p* < 0.001. **(C-D)** Quantification of **(C)** RNA expression and **(d)** tabular representation of average fold changes of *act5C*, *toy*, and *vglut* in tumor-derived neuroblasts (tNB) upon *med21* depletion. **(E-F)** Quantification of **(E)** RNA expression and **(F)** tabular representation of average fold changes of *act5C*, *toy*, and *vglut* in tumor-derived neuroblasts (tNB) and wildtype Neuroblas (NB). **(G)** Schematic representation of the mini-screen carried out to determine the role of Mediator in tumor growth. UAS-*BratRNAi;PointedGAL4* x *MedxRNA*i crosses were carried out in apple juice plates at 29°C. L1 larvae were collected every 3h to Vienna food plates to develop until L3 for 72hours at 29°C. L3 larvae were dissected and fixed brains were imaged (dorsal to ventral sides of the central brain) using confocal microscopy. Tumor volume quantification was performed using Imaris. **(H)** Tumor volume quantification relative to control for each of the indicated genotype. Genotypes are color-coded relative to the structural and functional modules those subunits belong to. ****p<0.0001; ***p<0.001; *p<0.05; **ns** – non-significant. **(I)** Z-projections of control (*BratRNAimcherryRNAi*), *BratRNAiMed14RNA*, and *BratRNAiMed21RNAi* tumors. Scale bar: 50µm.

To assess whether Mediator supports this elevated transcriptional activity in tumors, we measured the expression of previously identified hypertranscription-associated genes in FACS-isolated tNBs following *med21* knockdown. RT-qPCR revealed dramatic reductions in *act5C*, *toy*, and *vglut* expression (26.9-, 36.7-, and 45.2-fold, respectively; Figure 6C–D), demonstrating that Mediator is essential for sustaining hypertranscription in the tumor context. Importantly, expression of these same genes was significantly higher in tNBs compared to wild-type NBs (1.88-, 2.06-, and 1.24-fold increases for *act5C*, *toy*, and *vglut*, respectively; Figure 6E–F), validating the scRNAseq findings and confirming that Mediator-dependent transcription is further upregulated in tumors. We next asked whether Mediator-dependent hypertranscription is functionally required for tumor growth. To test this, we knocked down individual Mediator subunits in brat-driven tumors using the NBII-specific *pointed-GAL4* driver, co-expressing UAS-*myr*-GFP to enable tumor visualization. Under standardized rearing conditions, tumors were imaged and quantified for size (Figure 6G). Strikingly, depletion of multiple Mediator subunits resulted in a significant reduction in tumor size compared to control tumors expressing *mcherry* RNAi (Figure 6H).

Notably, all functional/structural modules have subunits whose depletion resulted in decreased tumor burden, including the kinase module. However, subunits from the head and middle modules were particularly important: nearly all knockdowns in these modules significantly reduced tumor burden, with the exception of *med11* (head) and *med1* (middle). In contrast, most tail module subunit knockdowns failed to reduce tumor size, suggesting either a lesser role in tumor progression or suboptimal RNAi efficiency. Among all tested subunits, depletion of *med14* and *med21* – previously shown to strongly impair hypertranscription – led to some of the most pronounced reductions in tumor size (Figure 6H-I), highlighting the importance of these subunits for sustaining the transcriptional state that supports tumor growth. Together, these results demonstrate that Mediator-dependent hypertranscription is a core requirement for tumor growth in *Drosophila* brain tumors. They position Mediator as a key regulator of the tumorigenic state in tNBs and support the broader relevance of hypertranscription as a conserved vulnerability in stem-cell-derived cancers.

## Discussion

We show that *Drosophila* NBs exist in a hypertranscriptional state relative to their progeny, characterized by broad gene expression, elevated Pol II occupancy, and activation of non-canonical NB genes. This state correlates with cellular potency and is reduced as cells differentiate into intermediate progenitors and neurons (Figure 7A). We identify the Mediator complex as a central regulator of hypertranscription: Mediator is broadly recruited across NB chromatin, and its depletion causes a strong, NB-specific reduction in transcription with minimal effects in differentiated cells, mirroring findings in mammalian systems where Mediator supports pluripotency and reprogramming but is dispensable for somatic cell survival(Li et al. 2015). The phenotypic spectrum resulting from subunit depletion is consistent with known Mediator architecture: loss of the core scaffold Med14 produces severe defects, highlighting its structural importance(Cevher et al. 2014), whereas depletion of tail or kinase module components, which are not essential for overall complex structural integrity(Jeronimo et al. 2016), results in little to no phenotype. The absence of phenotype after knock-down for some of Mediator subunits could also be due to redundancy or low RNAi efficiency. Although Mediator had not been previously directly linked to hypertranscription, inhibition of CDK8/19, the kinase module that inhibits Mediator activity, has been shown to sustain naïve ESCs, known to have a higher transcriptional output, and prevent their transition to the primed state(Lynch et al. 2020), suggesting a conserved role for Mediator in regulating transcriptional output and cell state across species.

**Figure 7.**
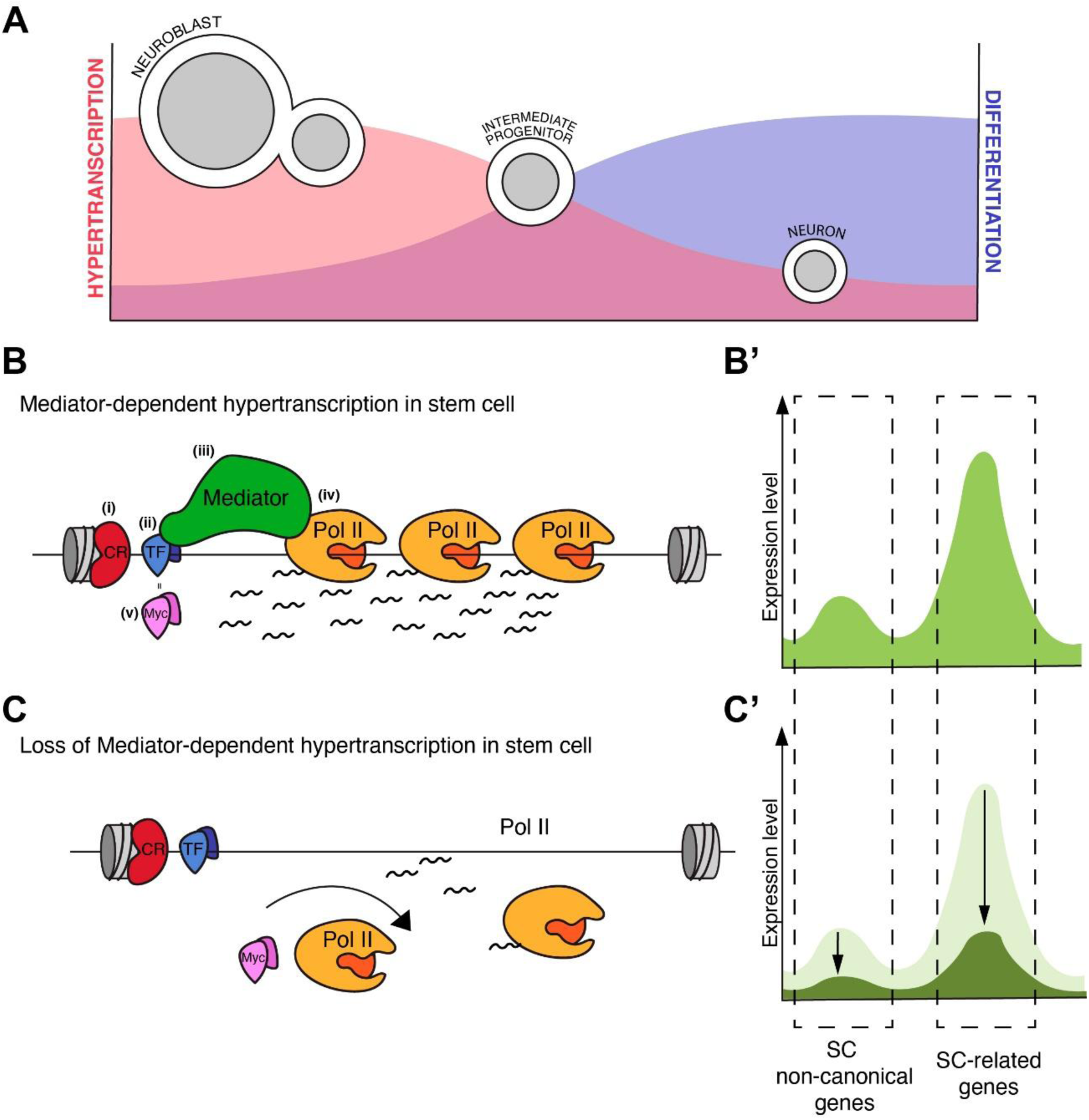
Mediator-dependent hypertranscription is required for stem cell identity and differentiation. **(A)** Hypertranscription correlates with the potency state of *Drosophila* neural lineages, with neuroblasts being more hypertranscriptionally active than more differentiated cells, such as neurons**. (B)** Schematic representation of the sequential molecular events underlying hypertranscription in neural stem cells. (i) Chromatin remodelers (CR) first establish a permissive chromatin landscape. (ii) Stem cell–specific transcription factors (TF), including pioneer factors, then engage enhancers or promoters to (iii) facilitate Mediator recruitment. (iv) Mediator binding promotes the assembly of general transcription factors and RNA polymerase II (Pol II), thereby initiating hypertranscription. (v) Myc, a TF whose expression is regulated by Mediator, subsequently amplifies the overall transcriptional output. **(B’)** As a consequence of hypertranscription, there is an overall increase in gene expression. This includes (i) neural stem cell–associated genes, such as transcriptional regulators, chromatin remodelers, and proliferation-related genes, which are expressed at higher levels and are essential for maintaining stem cell identity; and (ii) genes not canonically associated with stem cells, including muscle- or neuron-specific genes, which are expressed at lower levels and contribute to priming stem cells for subsequent differentiation. **(C)** Loss of Mediator ultimately leads to reduced recruitment of Pol II and consequent collapse of the hypertranscriptional state. **(C’)** An abrupt reduction in transcription compromises both stem cell identity and their capacity to differentiate. (i) Reduced expression of non-canonical stem cell genes may impair activation of lineage-specific transcriptional programs, thereby hindering differentiation. (ii) Concurrently, reduced expression of stem cell–associated genes disrupt core transcriptional networks and homeostatic processes, leading to defects such as prolonged or stalled cell cycles.

In recent years, efforts have been made to understand the molecular events sustaining hypertranscription. Chromatin accessibility is central to SC transcriptional capacity(Chen et al. 2024). Consistently, we find Pol II enriched at genes involved in chromatin organization in NBs but not neurons. This aligns with reports that link NB potency to chromatin accessibility(Aughey et al. 2018) and with the observation that chromatin remodelers such as Chd1 are essential for sustaining high transcription in mammalian cells(Gaspar-Maia et al. 2009). Moreover, chromatin remodelers have been shown to be essential for proper early mouse development(Stopka and Skoultchi 2003; Klochendler-Yeivin et al. 2000; Cao et al. 2003) and SC self-renewal, proliferation, and differentiation(Efroni et al. 2008; Xi and Xie 2005). These observations suggest that transcription-permissive chromatin enables hypertranscription. In ESCs, pioneer TFs such as Oct4 and Sox2 recruit remodelers to inaccessible chromatin(Soufi et al. 2012; King and Klose 2017) and act with Mediator to maintain SC transcriptional programs(Borggrefe and Yue 2011; Kagey et al. 2010; Malik and Roeder 2010b). Although these specific TFs are not conserved in *Drosophila*, NB-specific TFs likely play analogous roles. For example, Zelda, a NBII-specific pioneer factor acting in concert with Notch, binds extensively across enhancers to maintain NB identity(Larson et al. 2021). While a direct interaction between Zelda and Mediator remains untested, they may cooperate to sustain the hypertranscriptional state of NBs.

Through its ability to connect distal enhancers and TFs with Pol II and other general transcription factors (GTFs)(Schier and Taatjes 2020), Mediator serves as a critical hub in the transcriptional machinery, positioning both Mediator and GTFs as general regulators of transcription in NSCs. However, studies in *Drosophila* show that depletion of GTFs like TFIID and TBP only cause a small NB loss and mild transcriptional effects(Neves and Eisenman 2019), suggesting that GTFs primarily support basal transcription initiation rather than modulate transcriptional amplitude. In contrast, Mediator depletion produces a profound and NB-specific reduction of transcription, positioning Mediator as a selective and *bona fide* regulator of hypertranscription, distinguishing it from general transcriptional regulators that would be expected to uniformly disrupt transcription across lineages. Such a role is consistent with Mediator’s capacity to stimulate promoter-proximally paused Pol II release and favor productive elongation(Takahashi et al. 2011, 2015). Aligned with this model, naïve ESCs, which naturally operate in a more hypertranscribing state, exhibit reduced Pol II proximal pausing indexes, elevated elongation velocities, and increased RNA synthesis and turnover rates compared to primed ESCs(Shao et al. 2022), pointing to conserved principles that link Mediator activity, transcriptional dynamics and stem cell potency. Importantly, Myc, another known amplifier of already active genes(Nie et al. 2012), is bound by all analyzed Mediator subunits. However, *myc* knockdown phenotypes differ from those of Mediator depletion: while Myc perturbation variably leads to decreased NB proliferation or polarity defects(Song and Lu 2011; Rust et al. 2018), Mediator depletion consistently blocks differentiation and drives the emergence of ectopic NB-like cells. This suggests that Mediator establishes a transcriptionally permissive hypertranscriptional state that Myc later amplifies, in agreement with mammalian studies showing that Myc, together with YAP/TAZ, reinforces but does not initiate hypertranscriptional programs(Lavado et al. 2018b). Together, these findings support a model where chromatin accessibility and TF networks create a permissive state that Mediator translates into global hypertranscription, with Myc acting downstream to reinforce expression of active *loci* (Figure 7B and C).

A major open question in the field is whether hypertranscription simply marks stem cell identity or plays an active role in fate transitions. Our results show that hypertranscription is functionally required for NB differentiation. Loss of Mediator reduces hypertranscription and leads to the accumulation of stem-like cells, diminished neuronal output, and disrupted brain development. The reduced mitotic rate in ectopic NBs suggests delayed or stalled cell cycles. Interestingly, a previous study has shown that induction of hypertranscription via mutation of Topoisomerase I is sufficient to alter *in vitro* differentiation of mESCs into neurons(Lau et al. 2023). Similarly, another study has reported that *in vivo* exacerbation of hypertranscription via YAP/TAZ activation can cause neural progenitors to overproliferate and fail to differentiate(Lavado et al. 2018a). Thus, appropriate levels of hypertranscription are critical for both maintenance and timely exit from the stem cell state. ESCs are characterized by a more accessible chromatin, with regions of developmental regulatory genes being decorated with the counteracting histone marks H3K4me3 and H3K27me3. These bivalent chromatin regions, allow for very low-level transcription to prime for rapid activation of gene expression during lineage transitions(Kumar et al. 2021; Bernstein et al. 2006; Azuara et al. 2006). A similar mechanism may occur for hypertranscription, which may initiate expression of differentiation-associated genes at sub-threshold levels, ensuring they are poised for appropriate activation once cues signal the exit from the stem cell state (Figure 7B’ and C’). Importantly, the elevated transcriptional output of NBs cannot be attributed solely to differences in cell size, as NBs of markedly different sizes exhibit comparable numbers of detected gene features and RNA counts, and neurons with diverse volumes similarly display overlapping transcriptional outputs. Mediator depletion sharply reduces transcription in NBs without dramatically altering NB size, indicating that the decrease reflects a regulatory change rather than passive scaling. Consistent with this, several mammalian stem cell types, such as hematopoietic SCs, which are smaller than their differentiated progeny, also have elevated transcriptional output(Percharde et al. 2016).

We also examined whether metabolic alterations, classical regulators of stem cell function and potency(Bone et al. 2025), contribute to hypertranscription. In *Drosophila melanogaster*, tuning of the metabolic state by Mediator has been reported to be determinant to regulate terminal differentiation of NSCs during pupal stages(Homem et al. 2014). We found that while Mediator loss shifts metabolism toward glycolysis at the expense of OxPhos, hypertranscription itself remained remarkably resistant to metabolic perturbation. The observed metabolic shift may also partially reflect the reduced neuronal output of Mediator-depleted brains, as neurons largely rely on oxidative metabolism(Hall et al. 2012). Depleting glycolytic enzymes only modestly affected transcriptional output and produced no detectable lineage defects. These results indicate that although metabolism and transcription are tightly intertwined, hypertranscription is not simply a metabolic byproduct but reflects a separate regulatory layer essential for NB identity. In this regard, hypertranscription and metabolism may work in parallel to reinforce stemness.

Finally, our findings extend the significance of hypertranscription to tumor biology. Hypertranscription is increasingly recognized as a common feature of aggressive tumors(Zatzman et al. 2022; Cao et al. 2022). Our results reveal that *Drosophila* brain tumors co-opt the NB hypertranscriptional program to promote tumor progression. Tumor NBs display a strikingly enhanced transcriptional amplification, and Mediator depletion markedly reduces tumor size, demonstrating that hypertranscription is required to sustain tumor growth rather than an incidental consequence. Whether the molecular mechanisms underlying hypertranscription in tumorigenesis fully mirror those in physiological conditions remains to be determined. Interestingly, Myc is overexpressed in a majority of human cancers(Gabay et al. 2014) as well as in *Drosophila* brain tumors(Reichardt et al. 2018; Betschinger et al. 2006), and its loss is sufficient to reduce tumor burden in flies(Rebelo and Homem 2023). Given that dMyc is highly expressed and functionally important in both NBs and tNBs, it is likely that components of the molecular network regulating hypertranscription are preserved in both normal and tumor contexts. Taken together, our data support a model in which Mediator functions as a central node linking developmental transcriptional programs in NSCs with pathological transcriptional amplification in brain tumors, raising the possibility that targeting Mediator-dependent hypertranscription could represent a therapeutic strategy.

## Material & Methods

### Fly husbandry and stocks

Unless stated otherwise, fly crosses were set at 29°C and their progeny were maintained at 29°C to ensure maximum UAS/GAL4 expression and RNAi-mediated knockdown. The following RNAi lines were obtained from the Bloomington *Drosophila* Stock Center (BDSC): *UAS-med1^RNAi^* (34662), *UAS-med9^RNAi^* (33677), *UAS-med10^RNAi^* (34031), *UAS-med11^RNAi^* (34083), *UAS-med12^RNAi^* (34588), *UAS-med14^RNAi^* (34575), *UAS-med15^RNAi^* (32517), *UAS-med17^RNAi^* (34664), *UAS-med19^RNAi^* (33710), *UAS- med20^RNAi^* (34577), *UAS-med21^RNAi^* (34731), *UAS-med22^RNAi^* (34573), *UAS-med23^RNAi^* (34658), *UAS-med24^RNAi^* (33755), *UAS-med27^RNAi^* (34576), *UAS-med29^RNAi^* (37545), *UAS-cdk8^RNAi^* (35324), *UAS-cycC^RNAi^* (33753), *UAS-hexA*^RNAi^ (35155), *UAS-pfk*^RNAi^ (34336), and *UAS-pyk*^RNAi^ (35218). The following RNAi lines were obtained from the Vienna *Drosophila* Resource Center (VDRC): *UAS-med11^RNAi^* (106766), *UAS-med14^RNAi^* (105163), *UAS-med17^RNAi^* (105264), and *UAS-med21^RNAi^*. *UAS-mCherry^RNAi^* (BDSB#35785) or *w^1118^* (gift from António Jacinto) were used as controls depending on their adequacy to the type of experiment or due to convenience.

The following fly stocks were used to drive the expression of transgenes: ;;*VT201094-GAL4* (gift from Jürgen Knoblich), ;tubGal80^ts^;*VT201094-GAL4* (generated in the lab), ;;*VT201094-GAL4*,UAS-myr::GFP (generated in the lab), UAS-dicer2;*worniu-GAL4*, *asense-GAL80*; UAS-CD8::GFP (gift from Jürgen Knoblich), ;;*pointed*-*GAL4*, UAS-myr::GFP (generated in the lab), UAS-dicer2;*asense-GAL4*, UAS-CD8::GFP; (gift from Jürgen Knoblich), ;*earmuff-GAL4*, UAS-CD8::tomato; (gift from Jürgen Knoblich), UAS-dicer2;UAS-CD8::GFP;*earmuff-GAL4* (gift from Jürgen Knoblich), *nsyb-GAL4* (gift from César Mendes), and ;*UAS-brat^RNAi^*;*pointedGal4*,UAS-myr::GFP.

For the targeted DamID experiments, the following lines were used: ;;Dam (gifted by Andrea Brand), ;;Dam::PolII (gifted by Andrea Brand), ;;*UAS-Dam::Med11* (this study), ;;*UAS-Dam::Med17* (this study), ;;*UAS-Dam::Med21* (this study), and ;;*UAS-Dam::Med27* (this study). For the metabolites quantifications, the following lines were used: ;*worniu-GAL4*, *asense-GAL80*;UAS-Glucose sensor, ;*worniu-GAL4*, *asense-GAL80*;UAS-Laconic (original lines expressing the UAS-biosensor gifted by Stefanie Schirmeier).

### Generation of Dam-fusion transgenes

To generate pUAST attB-LT3-NDam-Med11, pUAST attB-LT3-NDam-Med17, pUASTattB-LT3-NDam-Med21, and pUASTattB-LT3-NDam-Med27, *Med11*, *Med17*, *Med21*, and *Med27* were PCR amplified from cDNA and cloned into pUASTattB-LT3-NDam (gifted by Andrea Brand) via Gibson assembly. Sequence confirmation was carried out by Sanger sequencing. Microinjection and transgenic animals generation was performed by The University of Cambridge Department of Genetics Fly Facility.

### Targeted DamID sequencing

Male transgenic flies expressing *UAS-Dam* (gifted by Andrea Brand), *UAS-Dam::PolII* (gifted by Andrea Brand), *UAS-Dam::Med11* (this study), *UAS-Dam::Med17* (this study), *UAS-Dam::Med21* (this study), and *UAS-Dam::Med27* (this study) were crossed to *tub-GAL80^ts^*;*VT201094-GAL4* female virgins. Egg laying was performed at 25°C for 6 hours and then raised at 18°C to prevent premature expression of Dam/Dam-fusion protein. Animals were transferred to 29°C 8 days after egg laying for 24 hours (corresponding to approximately wandering third instar larval stage at the time of dissection). One hundred brains were dissected and stored at −80°C until further use. Extraction of genomic DNA and enrichment/purification of methylated DNA was performed as previously described(Southall et al. 2013). Two independent biological replicates were generated for each experiment. Library preparations and sequencing were outsourced to Novogene (UK) Co., Ltd. DamID fragments were prepared for Illumina sequencing and all sequencing was performed as pair-end 150 bp using Illumina Novaseq 6000.

### Targeted DamID data processing

Next-generation sequencing reads in FASTQ format were aligned using the damidseq_pipeline software using default options for all data sets(Marshall and Brand 2015). The resulting gatc.bedgraph for each biological replicated of a single condition were averaged. Data sets were visualised using IGV. Genome browser tracks are represented as the average log_2_(Dam-fusion/Dam), where Dam-fusion is the protein of interest fused to Dam, while Dam is just the methylase and serves as background correction. Gene Ontology analysis was performed using the Database for Annotation, Visualization, and Integrated Discovery (DAVID) tools(Huang et al. 2009).

### Brain dissociation and cell sorting

Brains from wandering L3 larvae were dissected in Schneider’s medium (Sigma) and immediately transferred to supplement Schneider’s medium (10% fetal bovine serum (Sigma), 20mM glutamine (Sigma), 0.04mg/mL L-glutathione (Sigma), 0.02mg/mL insulin (sigma) Schneider’s medium (Sigma)). Afterwards, brains were enzymatically dissociated for 1 hour at 30°C in supplemented Schneider’s medium with 1mg/mL papin (Sigma) and 1mg/mL collagenase I (Sigma). After enzymatic dissociation, brains were washed twice with supplemented Schneider’s medium and mechanically dissociated in 200μL of supplemented Schneider’s medium using a pipette tip. The cell suspension was filtered through a 30μm mesh into a 5mL FACS tube (BD Falcon) and immediately sorted by fluorescence activated cell sorting (FACS) (FACS Aria III, BD). Neuroblasts (NBs), intermediate neuronal precursors (INP), and neurons were individually sorted according to (i) fluorescence intensity and (ii) size as described(Harzer et al. 2013). Purified cell populations were immediately collected in TRIzol LS (Invitrogen^TM^) to proceed to RNA extraction. When extraction did not take place immediately, samples were stored at −80°C until further use.

### RNA extraction, cDNA synthesis, and RT-PCR

RNA was isolated using TRIzol LS (Invitrogen^TM^) according to the manufacturer’s instructions. Extracted RNA was then treated with TURBO DNA-free^TM^ Kit (Invitrogen^TM^). cDNA was prepared using the RevertAid First Strand cDNA Synthesis Kit (Thermo Scientific^TM^). RT-PCRs were done using GoTaq qPCR Master Mix (Promega) on a QuantStudio^TM^ 5 Real-Time PCR System (Applied Biosystems^TM^). The following primers were used for amplification:

*act5C* Forward: GATAATGATGATGGTGTGCAGG;

*act5C* Reverse: AGTGGTGGAAGTTTGGAGTG

*toy* Forward: CTACAAGTGTCCAACGGTTGC

*toy* Reverse: GCTTTGAACCACCTATAGCTCG

*vglut* Forward: TGAGGTGCAATATGTCGGCG

*vglut* Reverse: TAGCCCCAGAAGAAGGACGA

*med21* Forward: TTTGAACGCATTGGACCCCA

*med21* Reverse: AGGTGTCGATGTCTTTAGCACA

To avoid normalization with housekeeping genes, expression of all genes was quantified in specific number of cells and relative levels were calculated using the ΔCt method(Huang et al. 2009). All measurements were done with technical triplicates.

### Immunofluorescence

Third-instar larval brains were dissected in ice-cold 1x phosphate-buffered saline (PBS), fixed in 4% paraformaldehyde-PBS for 30 minutes at room temperature and washed thrice in PBS with 0.0% Triton X-100 (PBT). Next fixed tissue samples were incubated for 1 hour in 1% Normal Goat Serum in PBT (blocking solution) and stained overnight at 4°C with primary antibodies. The next day, samples were washed in PBT and incubated for 30 minutes in blocking solution. Brains were then incubated for 2 hours at room temperature with secondary antibodies. During this incubation, samples were protected from light. Finally, brains were washed three times with PBT, incubated for 10 minutes with PBS, and mounted in Aqua Polymount (Polysciences, Inc). The following primary antibodies were used: guinea pig anti-deadpan (dpn, 1:1000, gift from Jürgen Knoblich), rabbit anti-asense (ase, 1:1000, in-house) or rat anti-asense (1:1000, gift from Jürgen Knoblich), rabbit anti-Dcp-1(1:250, Cell Signaling), mouse anti-pH3 (1:1000, Cell Signaling). The following secondary antibodies were used at 1:1000: goat anti-guinea pig Alexa^TM^ Fluor 568 (Thermo Fisher Scientific), goat anti-rabbit Alexa^TM^ Fluor 647 (Thermo Fisher Scientific), goat anti-rat Alexa^TM^ Fluor 647 (Thermo Fisher Scientific), goat anti-mouse Alexa^TM^ Fluor 405 (Thermo Fisher Scientific). Primary and secondary antibodies were diluted in blocking solution. Immunofluorescence images were acquired using a LSM880 (Carl Zeiss GmbH) confocal microscope. FIJI was used for image analysis.

### Quantification of NBs and their progeny

For the analysis of type II NB lineages, female virgins expressing UAS-dicer2;*worniu-GAL4*, *asense-GAL80*; UAS-CD8::GFP were crossed with *w^1118^* males (control) or males expressing RNAi against Mediator subunits. To analyze type I NB lineages, female virgins expressing UAS-dicer2;*asense-GAL4*, UAS-CD8::GFP; were crossed with *w^1118^*males (control) or males expressing RNAi against Mediator subunits. For quantification of number of NBs and their progeny in control or Mediator-depleted brains, immunofluorescent images of the posterior (NBII) or anterior (NBI) sides of brain lobes were acquired using LSM880 (Carl Zeiss GmbH) confocal microscope. NBIIs and INPs were identified using specific markers and classified accordingly: (i) NBII were the deadpan-positive cells located apically withing the lineage; (ii) immature INPs lack positivity for both asense and deadpan and are located adjacent to the NB; (iii) immature INPs asense-positive have staining for asense, but not deadpan; (iv) while mature are positive for both deadpan and asense. Only GFP-positive cells were considered and at least 4 distinct lineages from a minimum of 5 independent brains were quantified. For type I lineages, confocal images of the whole brain and NBIs (Dpn-positive, asense-positive cells) were counted and a minimum of 5 independent brains were quantified..

### Seahorse analysis

Respiration rates were measured in real-time using a Seahorse XF-24 analyzer (Seahorse Bioscience, North Billerica, MA). Animals expressing *UAS-mCherry^RNAi^*, *UAS-med17^RNAi^*, and *UAS-med21^RNAi^* were crossed with *VT201094-GAL4*, to induce the expression of the RNAi in central brain NBs. Third instar larvae of control and RNAi brains of the appropriate genotype were dissected in ice-cold Schneider’s medium (Sigma). The assay was performed in supplemented Schneider’s medium (10% fetal bovine serum (Sigma), 20mM glutamine (Sigma), 0.04mg/mL L-glutathione (Sigma), 0.02mg/mL insulin (sigma) Schneider’s medium (Sigma)). Following instrument calibration, brains were transferred to the XF-24 Flux analyzer to record basal cellular oxygen consumption rates (OCR) and extracellular acidification rates (ECAR) at a temperature of 29°C. One whole brain was used in a single well of a 24 well cell culture dish (Seahorse-bio). The single brain was placed in the bottom of each well, the organ was kept in place using a tissue screen supplied by the manufacturer, and 630uL of supplemented Schneider’s medium was added. The experiment was performed for 40 minutes according to the following details: 10 cycles, mix 1 minute, wait 0 minutes, and measure 3 minutes. At least 5 replicates were used for each genotype and 5 empty wells were included for background correction.

### Glucose sensor and Laconic Imaging

For glucose and lactate levels measurement in lineages *in vivo* in control (*UAS-mCherry^RNAi^*) and in brains with Mediator knockdown (*UAS-med21^RNAi^*), male flies expressing the RNAi were crossed with female virgins expressing either the UAS-Glucose sensor^86^ (gifted by Stefanie Schirmeier) or UAS-Laconic(Gándara et al. 2019b) (gifted by Stefanie Schirmeier). Expression of the biosensors and RNAi were driven in type II NBs by *worniu-GAL4, asense-GAL80*. Third instar larval brains were dissected in HL3-buffer (70 mM NaCl, 5 mM KCl, 20 mM MgCl2, 10 mM NaHCO3, 115 mM sucrose, 5mM trehalose, 5 mM HEPES; pH 7.35), mounted in a 35mm tissue culture imaging dish and immediately imaged using the sensitized emission approach with a Zeiss LSM980 confocal microscope (Zeiss). Images were acquired using a 40x water immersion objective and the following filter sets: excitation 405/emission 460/40 nm (CFP channel) and excitation 405/emission 520/50 nm (YFP, FRET channel). Z-projections of all slices were stacked into sum intensity using FIJI software. Then, a region of interest (ROI) containing all cells expressing the fluorescent sensor was defined and the mean gray value of all pixels was extracted for YFP and CFP. Ratios of the CFP/YFP were determined. Graphical representation and statistical analysis were made using the software GraphPad Prism 9 (GraphPad Software Inc., La Jolla, CA, USA). Unpaired two-tailed *t* test was used to determine statistical significance. Data is presented as mean ± standard deviation (SD). Statistical significancy was considered when *p*-values<0.05.

### Tumor Screen

Male flies expressing RNAi against individual Mediator subunits were crossed with *UAS-Brat-RNAi/CyoGal80;pointed-GAL4,UAS-myr-GFP* virgin females. The *pointed-GAL4* driver was used to specifically induce expression of both the RNAi constructs and the green fluorescent protein (GFP) in type II neuroblasts (NBs). *CyoGal80* was employed to suppress GAL4 activity, thereby preventing tumor formation via *Brat-RNAi* expression and allowing the maintenance of a stable fly stock.

All crosses were maintained at 29 °C to enhance the efficiency of the UAS-GAL4 system, ensuring robust expression of the RNAi constructs and GFP in the target cells. Crosses were initially set up on apple juice agar plates for 17 hours. Newly hatched first instar (L1) larvae were collected every 3 hours to ensure developmental synchronization and transferred to yeast-enriched food at 29°C. A maximum of 100 larvae were added to each plate to avoid overcrowding. Larvae were then kept at 29 °C for approximately 72 hours until reaching the third instar (L3) stage. For all experimental conditions, *UAS-mCherry-RNAi* was used as the control. After dissection, fixed brains were mounted in Aqua-Polymount (Polysciences, Inc.).

### Tumor volume quantification

Z-stacks of each brain were acquired using a Zeiss LSM 880 confocal microscope (Carl Zeiss GmbH) with a 1 µm interval between consecutive sections. Imaging was performed across the entire dorsal-to-ventral axis of both brain lobes to capture the full extent of the tumor. Tumor volume was quantified by generating a 3D reconstruction of the GFP signal from whole brains using Imaris software (Oxford Instruments). Representative images of tumor volume were generated as maximum intensity z-projections of all slices using FIJI (https://fiji.sc). Graphical representation and statistical analyses were performed using GraphPad Prism 9 (GraphPad Software Inc., La Jolla, CA, USA). Statistical significance was determined using an unpaired *t*-test. Data are presented as mean ± standard deviation (SD), and *p*-values < 0.05 were considered statistically significant.

## Data availability

Single-cell RNA sequencing data can be found in the Gene Expression Omnibus website (GSE179763). Published and newly-generated Targeted DamID data can be found in the Gene Expression Omnibus website (GSE77860; GSE282899).

## Acknowledgments

We thank the cytometry, microscopy, and fly facilities at NOVA Medical School for technical support and CONGENTO: consortium for genetically tractable organisms (LISBOA-01-0145-FEDER-022170), Bloomington *Drosophila* Stock Center (NIH P40OD018537) and Vienna *Drosophila* Resource Center (VDRC) for the stocks used in this study. We thank all members of the C.C.F.H. Lab for helpful discussions. We would also like to thank João Raimundo for discussion and reviewing the manuscript.

This project has received funding from the European Research Council (ERC) under the European Union’s Horizon 2020 research and innovation programme (H2020-ERC-2017-STG-GA 759853-StemCellHabitat to C.C.F.H.); by Wellcome Trust and Howard Hughes Medical Institute (HHMI-208581/Z/17/Z-Metabolic Reg SC fate to C.C.F.H.); European Molecular Biology Organization (EMBO) Installation grant (H2020-EMBO-3311/2017/G2017 to C.C.F.H.) and EMBO Long-Term Fellowship (EMBO-ALTF/1208-2018 to T.B.); by FCT (2023.16356.ICDT/LISBOA2030-FEDER 0070680 15940 – TUMORMETADYNAMICS and https://doi.org/10.54499/2024.17460.PEX. to C.C.F.H.; CEECIND/02989/2017 and EXPL/BIA-BID/1394/2021 to T.B.; 2020.05639.BD to A.R.R.). This work was also supported by iNOVA4Health (UIDB/04462/2020 and UIDP/04462/2020) and LS4FUTURE (LA/P/0087/2020) to C.C.F.H.

## Author Contributions

T.B. contributed to conceptualization, methodology, formal analysis, original draft writing, and funding acquisition. D.L. contributed to data acquisition and analysis. A.R.R. contributed to data acquisition and analysis. C.C.F.H. contributed to conceptualization, methodology, formal analysis, draft writing and reviewing, and funding acquisition.

